# Phenotypic Characterization of Liver Sinusoidal Endothelial Cells on the Human Liver-Chip for Potential *in vitro* Therapeutic Antibody Pharmacology Applications

**DOI:** 10.1101/2022.03.04.482954

**Authors:** Pelin L. Candarlioglu, Sushma Jadalannagari, Jake Chaff, James Velez, Sannidhi R Joshipura, Marianne Kanellias, Alexander P. Simpson, S. Jordan Kerns, Lorna Ewart, Manjunath Hegde, Jason Ekert

## Abstract

Liver plays a vital role in the human immune system, in the internalization and catabolic clearance of therapeutic antibodies and antibody-bound immune complexes via Fc-receptor (FcR) binding on the hepatic reticuloendothelial system cells. This Fc portion of the antibody binding to FcR in the liver initiates the clearance of these antibodies or immune complexes, which is vital in the context of half-life, dosing interval, efficacy, and safety of therapeutic antibodies. The liver sinusoidal endothelial cells (LSECs) express scavenging receptors that recognize, bind, and internalize an enormous diversity of extracellular ligands. The Fc gamma receptor FcγRIIB or CD32B on LSECs is responsible for the clearance of a large majority of IgG-bound immune complexes in the liver. Investigating the pharmacological effects of antibody clearance via human liver *in vitro* has been challenging due to the lack of reliable long-term LSEC culture protocols. Human LSECs downregulate the expression of CD32B rapidly *in vitro* in traditional 2D LSEC mono- and co-cultures,. We describe a Liver-Chip model with a co-culture of primary human LSECs and hepatocytes to recreate the liver microenvironment and extend the viability and function of LSECs, including CD32B expression levels, for a duration that is relevant for assessing the pharmacokinetics (PK) of therapeutic antibodies. Our results show that the expression of CD32B can differ based on experimental variables such as the source of primary cells (donor), passage number or source of detection antibodies used to visualize CD32B and shear stress. The CD32B expression was maintained for 14 days on the Liver-Chip in a donor-dependent but passage number independent manner. The Scanning Electron Microscopy (SEM) imaging showed the presence of fenestrae structures - one of the hallmarks of LSEC function. Key LSEC markers, including CD32B expression, were validated through flow cytometry.

## Introduction

Liver is a complex organ composed of an organized network of heterogeneous cells essential to maintaining homeostasis. The liver lobules are composed of parenchymal cells such as hepatocytes and numerous non-parenchymal cells (NPCs), including lymphocytes, antigen-presenting cells (APCs), stellate cells, and liver sinusoidal endothelial cells (LSECs)[1], [2].

The liver vasculature is an anatomically specialized structure where oxygenated arterial and nutrient-rich portal venous blood mix within a sinusoidal space that drains into a hypoxic central vein. The sinusoids form the fundamental functional unit of the liver and are lined with unique endothelial cells known as LSECs. Although their physiology is essential in key aspects of liver biology, the identification of LSECs is not straightforward. It requires tracking multiple cellular and phenotypic markers despite no standardized agreement within the field [3]. LSECs are typically distinguished from other endothelial cells by their reduced expression of CD34, CD31 (*PECAM1*), and von Willebrand factor (VWF) with elevated expression of LYVE1, CD14, CD32B (FcγRIIB), CD36, CD54, STAB1, STAB2, CLEC1B, and factor VIII (FVIII) (*F8*) [4], [5]as well as the presence of cellular structures such as transcellular fenestrations arranged in sieve plates [6], [7]. The native expression also changes based on the anatomical location within the liver as well as the distance from the central vein; Type 1 LSECs are classified as CD36^hi^CD32^-^CD14^-^LYVE1^-^CD54- and are furthest from the central vein, whereas Type 2 LSECs are LYVE-1^+^CD32^hi^CD14^+^CD36^mid-lo^CD54^+^ and located closer or adjacent to the central vein [8]. These distinct characteristics allow LSECs to carry out several critical functions within the liver, including the transport of metabolic blood components and drugs to hepatocytes through fenestrations, scavenging and clearing of antibody-antigen complexes from the blood and the secretion of the coagulation factor FVIII [9].

Among all the markers, FcγRIIB, a member of the FcγR family, is critical for the clearance of monoclonal antibodies (mAb) and plays a role in antibody-mediated liver toxicities [10]–[12]. CD32B is the sole inhibitory receptor among the family of FcγRs and binds to immune-complexed IgGs to modulate the immune response. It is expressed on a range of hematopoietic cells (e.g., B cells and myeloid cells) and endothelial cells such as the villus interstitium of the placenta and LSECs. It is estimated that up to 75% of all CD32B expressed in the body are on LSECs [13], [14]. Click or tap here to enter text.Click or tap here to enter text.CD32B also plays a pivotal role in removing small immune complexes such as IgG antigens or -opsonized viruses. In mAb therapy, CD32B does not only reduce the efficacy of some mAbs through its inhibitory function but is also essential for the action of immunomodulatory mAbs. CD32B can elicit a unique cross-linking function of immunomodulatory mAbs through a scaffolding effect to drive strong agonistic signals in the target cell [11], [12], [15], [16].

Over the last three decades, mAbs have emerged as a major class of human therapeutics on the market. Non-human primate (NHP) *in vivo* models are relied on to predict pharmacokinetics, immune-complex clearance, immunogenicity and human toxicities that are not always clinically translatable [17]. Assessing therapeutic antibody pharmacokinetics and pharmacodynamics (PKPD) in NHP models sometimes exaggerate antibody-dependent autoimmune and hypersensitivity responses and vasculotoxicities observed in humans, such as with agonistic mAbs directed against CD137/4-1BB [18], GSK305002 [19] and other immunomodulatory mAbs [12], [20], [21]. These toxicities were mediated by LSECs, which are often overlooked in assessing unexpected toxicities of therapeutic mAbs. Liver toxicity tests historically focus on hepatocyte-mediated PK and toxicities of small molecule-based drugs.

There is a need for better prediction of mAb toxicities and clearance in therapeutic antibody discovery before ‘first-in-human’ use. Pre-clinically, human liver models have primarily focused on monocultures of hepatic cell lines such as HepG2 and HepaRG or primary human hepatocytes in 2D and 3D formats. Co-culture iPSC [22]-, organoid [23] - or microphysiological systems-based models [24]–[26] that include hepatocytes and non-parenchymal cells focus on maintaining hepatocyte function. This approach is relevant to recapitulating hepatocyte-mediated absorption, distribution, metabolism, elimination, and toxicity (ADMET) of small molecule-based drugs or hepatocyte-related disease conditions such as non-alcoholic steatohepatitis (NASH) [27]. Primary LSECs grown at 2D under static *in vitro* conditions rapidly lose the expression of critical phenotypic markers such as CD32B and the presence of fenestrae [28]–[30], leading to a functional decrease in mimicking hepatic clearance of antibody complexes.

Because of these challenges [31] with the current *in vitro* LSEC models and the limitations of NHP-based *in vivo* models in predicting the PKPD of therapeutic antibody candidates [17], we developed a primary human LSEC-hepatocyte co-culture model using Emulate’s commercially available Liver-Chip model. This Liver-Chip model has previously focused on the characterization of hepatocyte functionality and hepatocyte mediated liver toxicity but not on LSEC viability and functional phenotypes [25]. Here we extensively characterized the viability and function of LSECs, including CD32B expression levels, for 14 days, relevant for assessing the PK of therapeutic mAb candidates. We evaluated the effect on the co-culture of LSECs and hepatocytes from four LSEC donors with four passage numbers, two CD32B different detection antibodies, and 0.07dyne/cm^2^ shear stress as the mechanical stimuli to promote CD32B expression. Our results show that the expression of CD32B can differ based on experimental variables such as the source of primary cells (donor), passage number, or source of detection antibodies used to visualize CD32B. The CD32B expression was maintained for 14 days on the Liver-Chip in a donor-dependent but passage number-independent manner. SEM imaging showed the presence of fenestrae structures as one of the hallmarks of LSEC morphology, and the key LSEC markers, including CD32B expression, were validated using flow cytometry. We believe this model presents an avenue for the safety assessment of immuno-modulation of biologics in the liver and the potential to predict the pharmacokinetics/pharmacodynamics of therapeutic antibodies in humans.

## Methods

### Human Co-Culture Liver Chip Fabrication and Cell Culture

Four independent donors of primary human LSECs (Cell Systems and Upcyte) were thawed and cultured according to vendor protocols. Donor one was obtained from Upcyte (Donor 462), while donors two and three were from Cell Systems (566.01.01.01.1Tb and 566.02.03.05.0M, respectively). Donor four is iXCell (200356). Upcyte cells were chosen for this study as they are genetically engineered to induce transient proliferation capacity like the native liver [23]

45 Liver-Chips were used in this study, 15 per donor. Of these, 4 were terminated on Day 3 and 4 on Day 7 post-LSEC seeding with the remaining Liver-Chips terminated on Day 14 post LSEC seeding. 3 Liver-Chips were either fixed for immunofluorescence imaging, SEM imaging or stained and analyzed for marker expression by Flow Cytometry. Donors 2 and 3 were evaluated in two independent experiments.

The Liver-chip coating, seeding, and connection were conducted according to previously published protocols[24], [25]. Briefly, before cell seeding, Liver-Chips (S1 Chip^®^, Emulate Inc.) were surface-functionalized using reagents ER-1^™^ (Emulate reagent: 10461) and ER-2^™^ (Emulate reagent: 10462). The Liver-Chips were coated with extracellular matrix (ECM) components; 100 μg/mL rat tail collagen type I (Corning^®^) and 25 μg/mL bovine fibronectin (ThermoFisher) in dPBS (Sigma). Primary human hepatocytes (Gibco) were seeded in the top channel at a density of 3.5 x 10^6^ cells/mL using complete hepatocyte seeding media. After 24 hours, to promote a three-dimensional sandwich culture, the hepatocytes were overlayed with 200 μL hepatocyte maintenance media with 2.5% Matrigel^®^ (Corning^®^).

On the day of LSEC seeding (Day 0), donor one and donor three LSECs were harvested from the flasks by adding 3 mL trypsin (Sigma) for 2-3 minutes at 37°C to detach the cells from the monolayer. Trypsinization was stopped by adding complete LSEC media with 10% FBS. Cells were then counted and seeded into the bottom channel of the Liver-Chip at a density of 3 x 10^6^ cells/mL. Donors two and four LSECs were thawed from the vial and seeded directly onto the Liver-Chip at the same density. Four hours later, all the Liver-Chips were washed with respective media (hepatocyte maintenance media for top channel and LSEC complete media for bottom channel) and connected to Pods^®^ with 4 mL respective media in the reservoirs. Media was perfused in a Zoë^®^ perfusion system and equilibrated according to standard Emulate protocols.

The bottom channel flow rate was set to 60 μL/h while the top channel flow rate was set to 30 μL/h. The bottom channel flow rate was increased to 250 μL/h after 24h, while the top channel flow rate remained at 30 μL/h to maintain shear stress of 0.07 dyne/cm^2^. Once flow had started, media was refreshed daily in the bottom channel at 8 hours and 16 hours, while for the top channel, media was refreshed every 48 hours.

#### Biochemical assays

Effluent from the outlet of the top channel was used to quantify albumin levels on Days 3, 7, and 14 post LSEC seeding (n=4 per time-point per donor). A sandwich albumin ELISA (Abcam, ab179887) was run in accordance with the supplier’s protocols. Hamilton Vantage liquid handling platform performed all steps prior to the 1-hour incubation. Effluent samples were diluted in a 1:500 ratio and the absorbance was measured with the Synergy Neo Microplate Reader (BioTek) at 450 nm.

#### Morphology and Immunofluorescence staining

For morphological assessment on Days 3, 7 and 14 post LSEC seeding, 4 to 6 brightfield images were acquired for each Liver-Chip (n=4 per donor). The brightfield images were acquired on an ECHO Revolve microscope with a 10X objective.

On Days 3, 7 and 14 post-LSEC seeding, Liver-Chips were detached from Pods^®^ and washed once with 200 μL 1X PBS. Both channels (n=3 per donor) were fixed using 4% paraformaldehyde (PFA) solution (Electron Microscopy Sciences). The PFA solution was pipetted into both channels followed by a 15-minute incubation at room temperature (RT). The Liver-Chips were washed with 1X PBS and stored at 4 □C until immunofluorescent staining. Liver-Chips were permeabilized with 0.125% Triton^™^ X-100 (Sigma) for 10 minutes at room temperature then washed with 200 μL 1X PBS (Sigma). Samples were then blocked with 2%BSA (Sigma), 10% normal goat serum (Life Technologies) blocking solution for 1 hour at RT (100 μL/channel). Next, primary antibodies were prepared at 1:100 in a 1:4 blocking solution (2% BSA (Sigma), 10% normal goat serum (Life Technologies) diluted in 1X PBS. The top channel of each Liver-Chip was left in 1X PBS. The bottom channel of each Liver-Chip was stained with rabbit anti-Stabilin-1 (Novus Biologicals), mouse anti-LYVE-1 (ThermoFisher) and FITC conjugated CD32B AT10 clone (University of Southampton) antibodies. Then, 100 μL of this primary staining solution was added to the bbottom channel of each Liver-Chip and was incubated at 4°C overnight. The Liver-Chips were then washed twice with 1X PBS. Secondary antibodies were prepared at 1:1000 in a 1:4 blocking solution diluted in 1X PBS. The bottom channel of each Liver-Chip was stained with donkey anti-mouse AF555 (ThermoFisher) and donkey anti-Rb AF647 (ThermoFisher). Once added, Liver-Chips were incubated at RT in the dark for 2 hours, washed twice with 1X PBS, stained with DAPI (ThermoFisher) 1:1000 for 10 minutes at RT, and washed once more with 1X PBS. The stained Liver-Chips were stored in 1X PBS until imaging.

#### Image Acquisition

Fluorescent confocal image acquisition was performed using the Opera Phenix^™^ Plus High-Content Screening System and processed in Harmony 4.9 Imaging and Analysis Software (PerkinElmer). Liver-Chips were placed in the Emulate 12-Chip Imaging Adapter (Emulate 21008-01). The following parameters were used for image acquisition: 20X Water N.A. 1.0 objective, DAPI (Time: 180ms, Power: 80%), TRITC (Time: 200ms, Power: 80%), Alexa 488 (Time: 200ms, Power: 80%) and Alexa 647 (Time: 200ms, Power: 80%). Z-stacks were generated with 5 μm between slices for 21 planes. Three fields of view (FOVs) per Liver-Chip were acquired, with a 5% overlap between adjacent FOVs.

#### Scanning Electron Microscopy (SEM) Analysis

SEM was performed at the Harvard Medical School Electron Microscopy Facility with samples from Days 3, 7 and 14. One Liver-Chip per donor was selected, and the channels were washed with PBS and fixed in 2% glutaraldehyde in 0.1 M sodium cacodylate buffer (Electron Microscopy Studies) of pH 7.4, at 4°C overnight. The PDMS from the bottom of the Liver-Chips was cut and peeled to expose the bottom channel for SEM imaging. Further, the centers of the Chips were cut into 15 mm diameter pieces for easy mounting on the SEM imaging stubs. After overnight fixation at 4°C, the cut Chips were rinsed with 0.1 M sodium cacodylate buffer and fixed in 1% osmium tetroxide in 0.1M sodium cacodylate buffer at 37°C for 1 hour. The cut Chips were then rinsed in water three times for 5 minutes each and dehydrated using 30% ethanol for 5 minutes, 50% ethanol for 5 minutes, 70% ethanol for 5 minutes, 95% ethanol for 10 minutes, and 100% ethanol for 10 minutes. Specimens were then mounted onto stubs using conductive carbon tape, silver-painted using colloidal graphite, dried overnight, and sputter coated. Samples were then dried using HMDS protocol in 100% ethanol for 1 hour, 100% HMDS twice for 30 minutes, and air-dried overnight in the hood. Images were obtained using Hitachi S-4700 SEM at 3KV. For every Chip, five different images were obtained from different FOVs. Images were at a magnification of 1μm. Fenestrations were measured in ImageJ using the scale bar provided on each image to set the scale, and a line was drawn across the diameter of each observed fenestration.

#### FACS Analysis

LSECs were harvested from the Liver-Chip on Day 14 using PBS-EDTA (ThermoFisher). Briefly, the Liver-Chips were disconnected from the Pods^™^ and both channels were washed once with PBS to remove the media. While the top channel was blocked by pipette tips with PBS, the bottom channel was incubated with 200 μL PBS-EDTA for 10 minutes. The LSECs were triturated by pipetting up-and-down several times, and cells were collected in 96-well v-bottom plates (Falcon). All the cells were stained for live/dead cells using ZombieAqua^™^ viability dye (Biolegend) at 1:100 dilution, according to the manufacturer’s instructions. The harvested cells were live stained for the cell surface antigens using the following antibodies: FITC anti-human CD32B (AT10-clone, University of Southampton), Brilliant Violent 421 antihuman CD54 (Clone HA58, Biolegend 353132), and Brilliant Violent 785 anti-human CD14 (Clone 63D3, Biolegend 367142) in Cell Staining Buffer (Biolegend) at 1:25 dilution. Stains were added to the appropriate wells with cells with fluorescence minus one (FMO) and single stain wells and incubated at RT for 30 minutes. After staining, cells were washed and resuspended in staining buffer. Data were acquired using BD FACS Celesta^™^ flow cytometer (BD Biosciences) and was analyzed with FlowJo V10 software (FlowJo).

## Results and Discussion

### Human Liver-Chip maintains the morphology of LSECs for 14 days of culture

Traditional cell culture models such as monocultures of hepatic cells in 2D or 3D formats or co-culture with other NPCs have limited capacity to recapitulate physiological microenvironments of the human liver. These models also often focus on hepatocyte function and morphology. In addition, LSECs are often regarded as a poorly characterized cell type that can rapidly dedifferentiate and lose expression of functional phenotypic markers such as CD32B *in vitro* [5]. Hence, we show for the first time that the Liver-Chip was able to promote and maintain classic endothelial cell morphology for four independent LSEC donors (Figure 1b, Donor 4 in Supplementary figure 1a). Qualitatively, all the donors maintained prototypical endothelial morphology for 14 days. However, donors 2 and 4 demonstrated approximately a 20% decline in cell number by Day 14 as observed through bright-field imaging (scoring performed according to supplementary figure S2, donor 4 in supplementary figure S3). Similar LSEC morphology was demonstrated by donor one in a liver organoid model [23] and by donor three in human coculture [25], [30] and quad-culture [24] Liver-Chips, respectively.

**Fig. 1.**
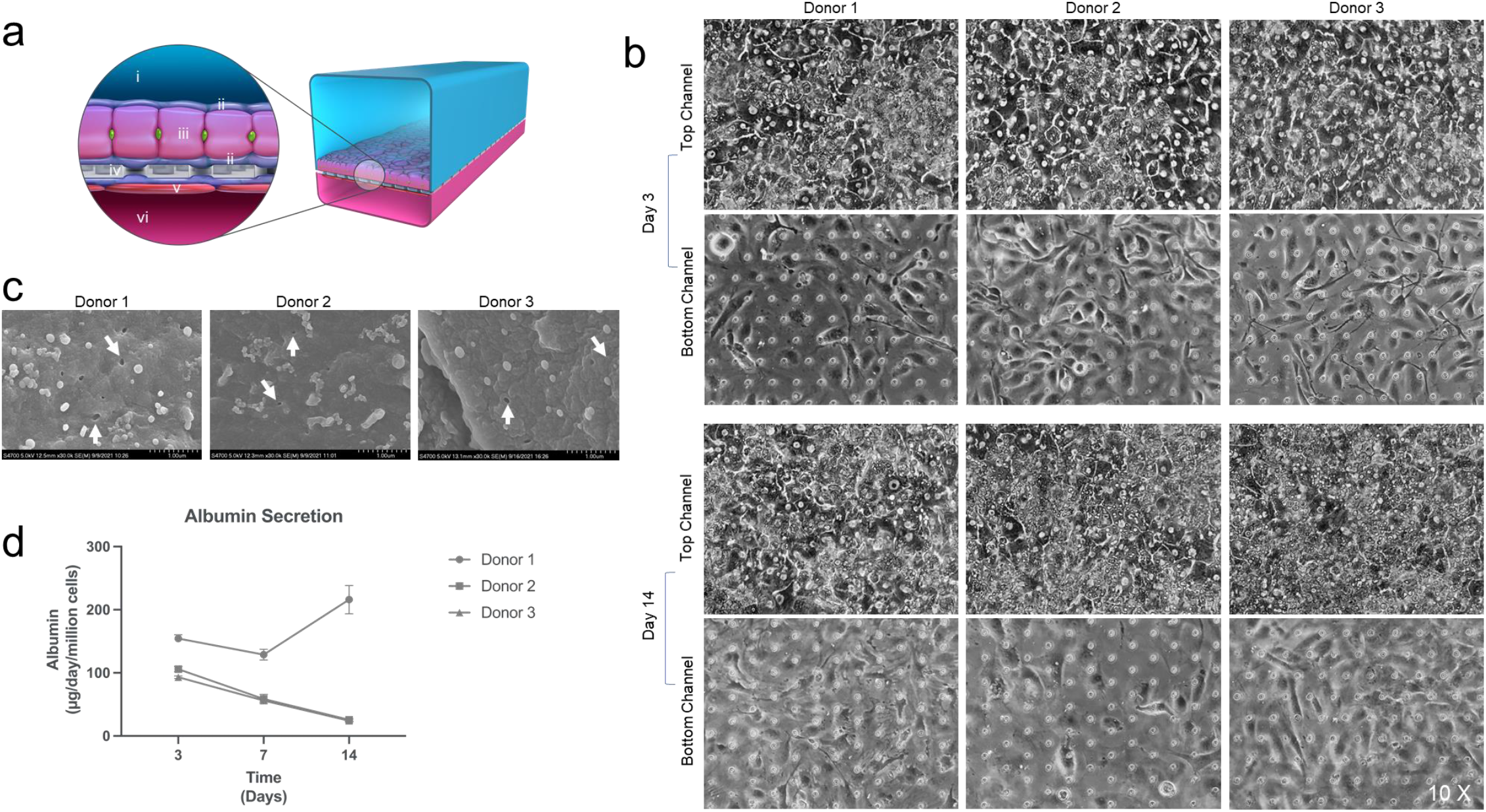
Assessment of cellular morphology, function, and toxicity on the Liver-Chip. a) The Liver-Chip has two channels, a top channel (i) and bottom channel (vi) separated by a porous membrane (iv). Hepatocytes (iii) are seeded onto the top channel and overlayed by Extra Cellular Matrix (ECM) components collagen and fibronectin to create a sandwich culture (ii) LSECs (v) are seeded in the bottom channel. b) Hepatocyte and LSEC morphology of Donors 1, 2, and 3 (from left to right) showed classic hepatic and endothelial morphology on Day 3 (top panel) and Day 14 (bottom panel). Increased debris accumulation was seen on Day 14 for all hepatocytes. A reduction in cell numbers was seen for donor 2 LSECs by Day 14. The arrangements of circles seen in the images are membrane pores. c) SEM images of donors one, two, and three (from left to right) were taken to verify fenestrations on the cells’ surface. White arrows indicate the presence of sparsely distributed perforations and small fenestrations. d) Albumin secretion was measured on Days 3, 7, and 14 from the Chip top channel effluents. Donors 2 and 3 had a steady decrease in albumin production by Day 14 while donor 1 increased by Day 14.

SEM is the gold standard for visualizing the ultrastructure of cells and reveals that 2-20% of the LSEC surface is covered with fenestrae that are either individually scattered or arranged in a sieve-like pattern [32] with no basement membrane, and no cytoplasmic projections such as filopodia [33]. Fenestrations are characteristic transcellular pores that allow for the uptake of small molecules from the blood to the surrounding hepatocytes and vice versa [34]. However, visualizing LSEC fenestrae through SEM is historically difficult due to the loss of fenestrations within hours of isolation and culture [35]. SEM imaging of LSECs co-cultured with hepatocytes on Liver-Chip identified sparsely distributed perforations and small fenestrations of ~100 to 200 nm in donors 1,2, and 3, demonstrating LSECs can be cultured and maintained with fenestrations on the Liver-Chip (Figure 1c). Similar fenestrations of 100-200 nm were observed when LSECs were cultured for 1 day on collagen I coated glass on a different liver sinusoid chip model[36].

### Effect of LSECs on hepatocyte albumin production

Primary human hepatocytes cultured in 2D rapidly lose physiological function as indicated by a decrease in albumin production [37]. Hence, to evaluate the function of hepatocytes on Liver-chips, albumin levels were measured from the Liver-Chips effluent of the top channel. The hepatocytes on Liver-Chips maintained physiologically relevant albumin levels [25], [38] over time, although hepatocytes cultured with donor one LSECs demonstrated higher albumin levels (150 – 200 μg/day/million cells on Days 3 to 14) in comparison to the other LSEC donors 2 and 3 (~100 to 25 μg/day/million cells on Days 3 and 14 respectively) (Figure 1d). Albumin levels are expected to decrease in parallel with a decline in cell health (as seen for donor 2 and 3). Interestingly donor one demonstrated a sustained elevated level of albumin (physiologically relevant *in vivo* albumin levels are 20 to 105 μg/million/day) over the experimental period [24], thus showing LSECs affect the functionality of hepatocytes in culture. We hypothesize this to be due to vascular endothelial growth factor (VEGF) secreted by the endothelial cells, which is known to mediate hepatocyte growth by paracrine signaling [39].

### Liver-Chip maintains immunophenotypic marker expression in LSECs for 14 days of culture

After confirming relevant albumin levels, we utilized CD32B as a marker to distinguish LSECs from other liver cell types and is involved in the regulation and uptake of antibodies and other immune complexes [5]. A model that maintains CD32B expression levels is crucial to understanding LSECs and their role in uptake and clearance [14]. Thus, to evaluate the phenotypic marker expression of LSECs on human Liver-Chip, we used immunofluorescence imaging to compare cell surface markers such as Stabilin-1, LYVE-1, and CD32B (Figure 2a-c). All the donors qualitatively demonstrated consistent expression of LYVE-1 and Stabilin–1 for 14 days through immunofluorescence staining. Donor one demonstrated CD32B expression only on Days 7 and 14, while donors two and three showed low but consistent expression throughout the experiment. Thus, the LSECs used in this study are isolated from zones 2 and 3 of the acinar liver lobule. However, donor 4 demonstrated stabilin-1 expression but were negative for CD32B and LYVE-1 over time, indicating these LSECs were isolated from zone 1 of the liver lobule (Data in supplementary figure 1b).

**Figure 2:**
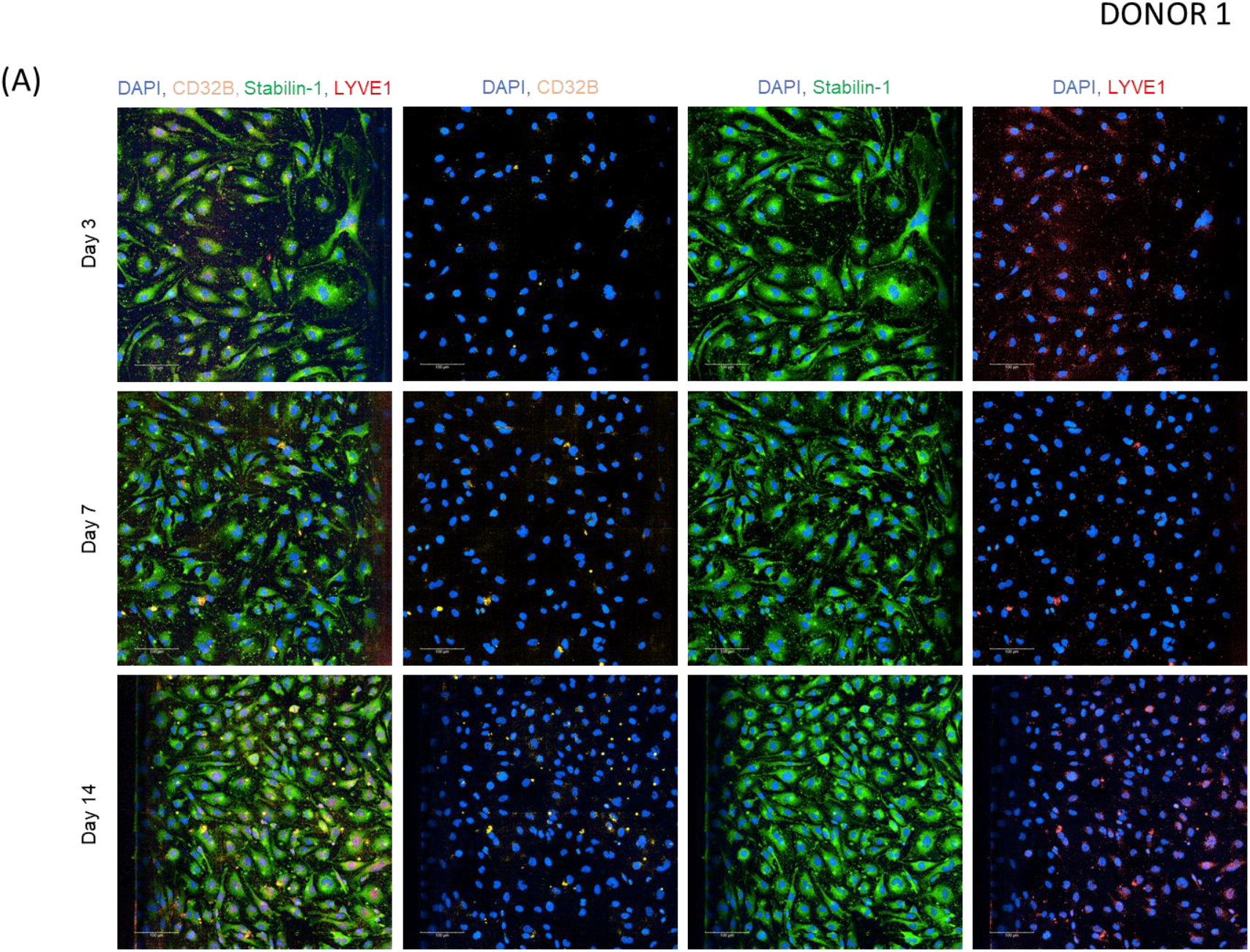
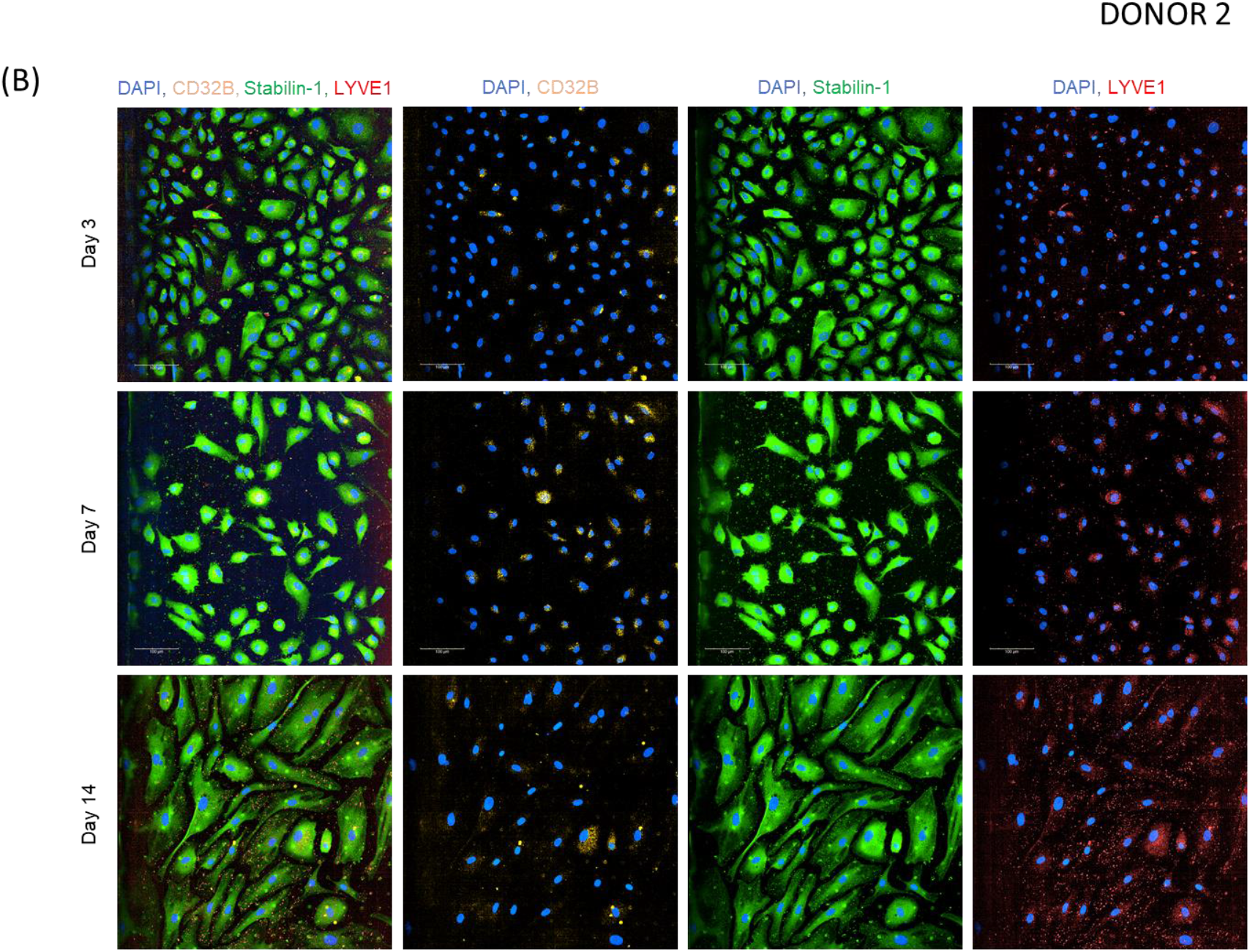
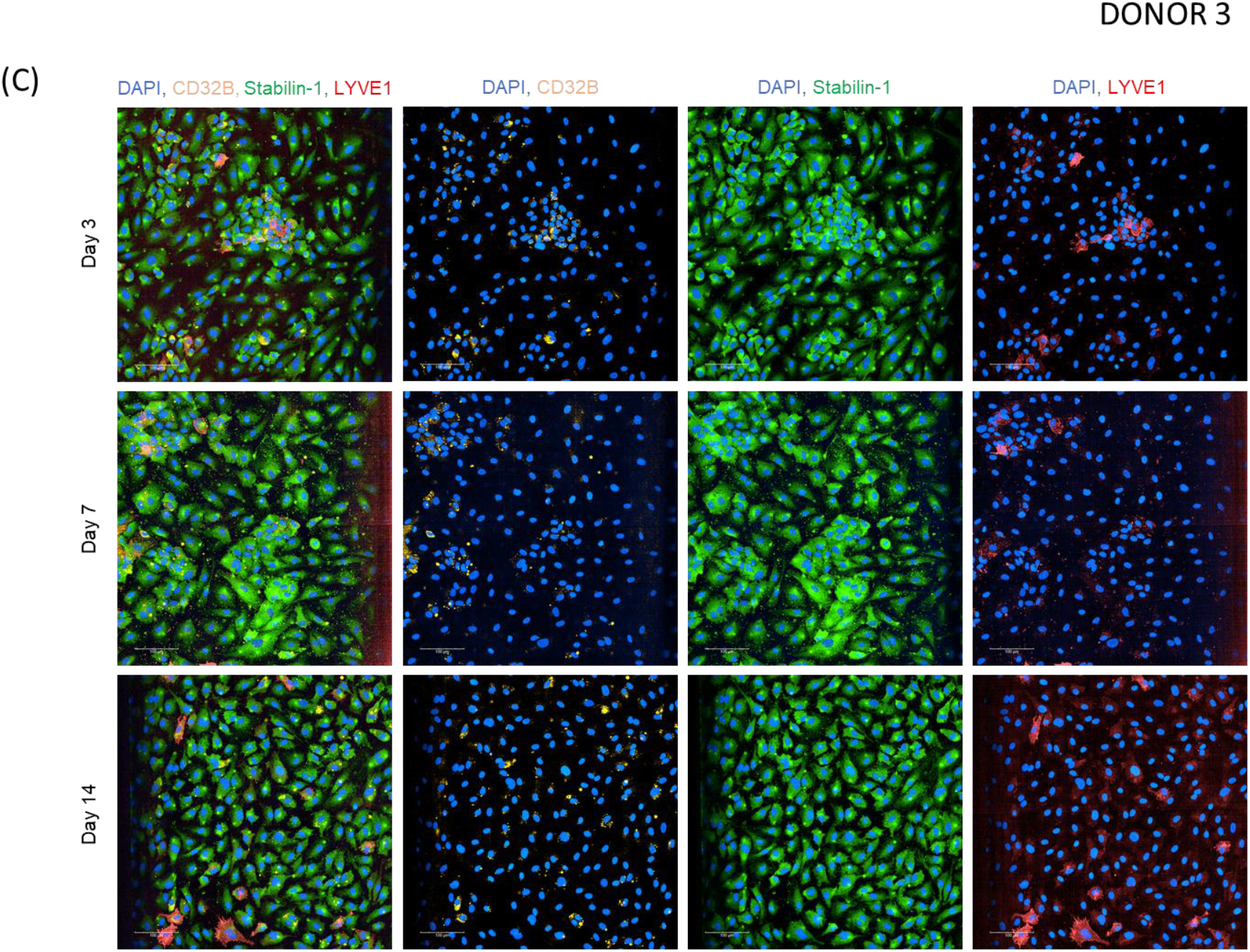
Representative immunofluorescence microscopy showing CD32B, Stabilin-1, LYVE-1 immunofluorescent staining and DAPI nuclear staining in LSECs from three different donors. From left to right composite images and their respective split-channel overlays with DAPI can be seen. Black edges indicate the edge of the main Chip channel. On Days 3, 7 and 14, three Chips from each donor group were fixed and stained for LSEC markers such as CD32B, Stabilin-1, and LYVE-1. All donors were Stabilin-1^+^ LYVE-1^+^ from Day 3 to 14, indicative of type II LSECs. A subpopulation of a) donor one cells was CD32B^+^ by Day 7, b) donor two cells are CD32B^+^ by Day 3, however, there was a decline in cell numbers by Day 14 in comparison to the other donors, c) a small population of donor three cells were also CD32B^+^ from Days 3 to 14 of culture on Liver-Chip.

Furthermore, donor one LSECs have previously been shown to maintain strong expression of CD32B and vascular endothelial cadherin (VE-Cadherin) in an *in vitro* organoid model of the liver sinusoid. These cells were co-cultured with tissue-resident macrophages on a Dynamic42 biochip and perfused at a shear rate of 0.7 dyne/cm^2^ [23],[40]. However, channel the Liver-Chips used in this study were maintained at a shear rate of 0.07 dyne/cm^2^ due to the differences in channel dimensions. In both conditions, the donor 1 LSECs demonstrated similar CD32B expression on Days 7 and 14. CD32B expression was also evaluated in donors 2 and 3 using two different clones of CD32B antibody (AT10 FITC and polyclonal against mouse myeloma cell line NS0-derived recombinant human Fc gamma RII/CD32 Ala46-Pro217, supplementary figure S4), with both showing expression of CD32B. It is important to note AT10 and other polyclonal antibodies are not CD32B specific. However, LSECs have been extensively characterized and shown to only express CD32B, hence it is assumed in the context of this work to be CD32B. Since LSECs express up to 90% of CD32B present in the liver[13], a substantial portion of immune-complex elimination is accounted for by LSECs. Hence, future studies with LSECs on Liver-Chip can be designed to target the rate of CD32B uptake for different immune complexes, allowing for a deeper look into LSEC function.

Additionally, to further characterize LSECs on the Liver-Chip, cells were harvested from the bottom channel on Day 14 and characterized by flow cytometry for the expression of endothelial markers including CD54, CD14, and CD32B (Figure 3b). LSECs from the Liver-Chip were also assessed for Stabilin and LYVE-1 expression by flow cytometry, however, no expression was detected by this method (data not shown). This could be because the method used for isolating cells could have damaged these markers. Fomin and colleagues demonstrated that in freshly isolated fetal liver cells, nucleated hematopoietic cells expressed high CD45 and low CD14, hepatoblasts had high CD326 and low CD14, while LSECs had high CD14 expression and no CD45[18]. Similarly, Karrar and colleagues compared unstimulated and stimulated LSECs and human aortic endothelial cells and reported that only LSECs demonstrate a high expression of CD14 and CD54 [42]. Hence, when LSECs were isolated from the Liver-chip and gated to select only cells negative for the viability dye AquaZombie^™^, all 3 donor LSECs on the Liver-Chip demonstrated a marked increase in CD54 expression, with donor three expressing the highest levels of CD54. To ascertain the percentage of positive cells, the shift in fluorescence signal was compared to the FMO to identify cells expressing the receptors. All donors exhibited similar percentage CD14 and CD32B expression, thus proving the Liver-Chip maintains the characteristic LSEC phenotype and marker expression for 14 days. Additionally, when the expression of these markers was compared to cells directly from the vial for donor three (passage 3), LSECs on Liver-Chip showed an upregulation of CD54, and a modest increase in CD14 and CD32B, thus indicating the co-culture of hepatocytes and LSECs under flow on the Liver-Chip improved the phenotypic marker expression of LSECs, which more closely emulates *in vivo* liver (Figure 3c). Cells directly from the vial were not compared for donors 1 and 2 due to low cell counts.

**Fig. 3.**
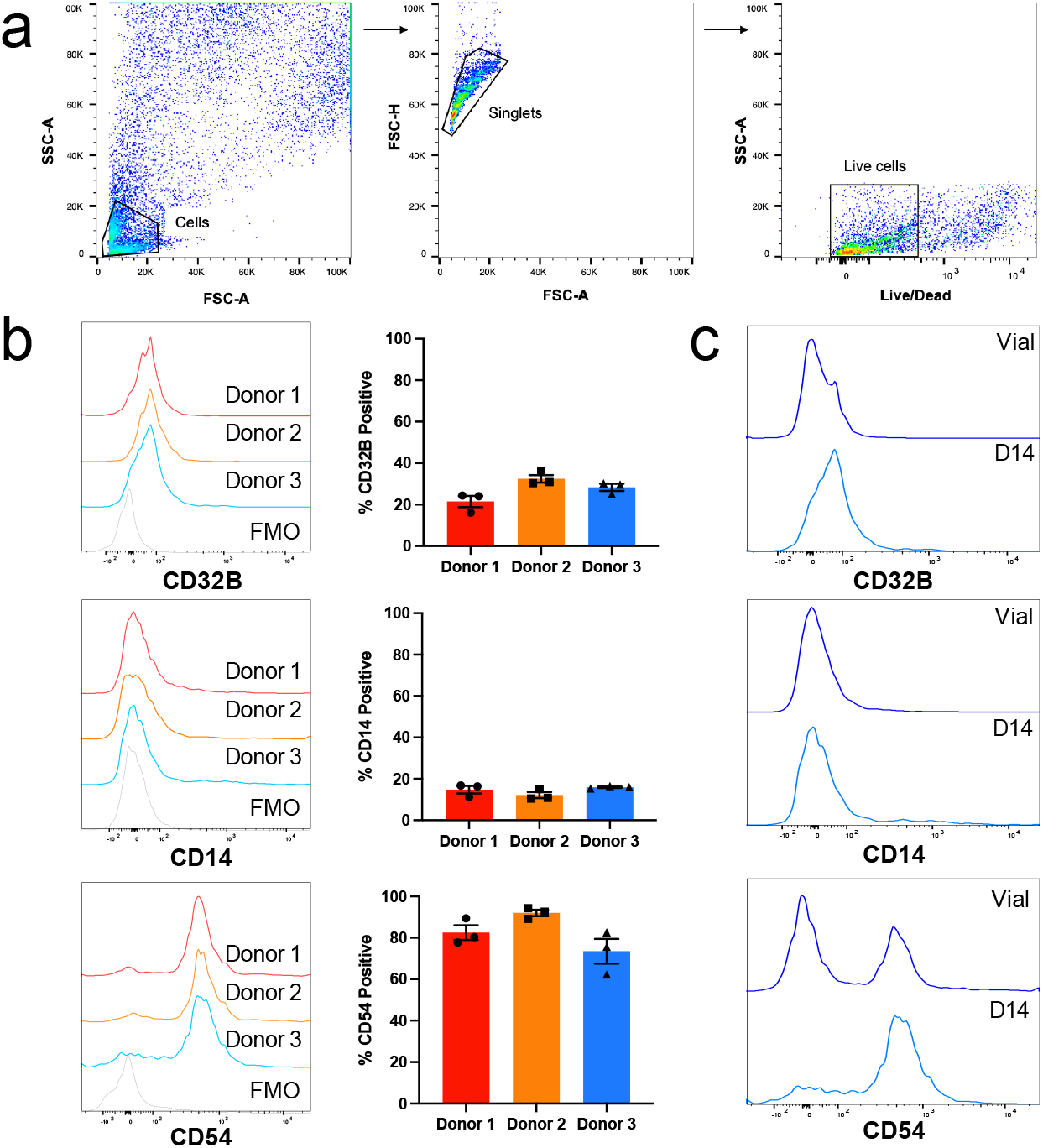
Expression of cell markers CD32B, CD14, and CD54 <u>at Day 14</u> by flow cytometry. a) Sequential gating strategy showing the selected cell population followed by gating for Live/Dead stain, CD32B, CD14, and CD54. In the leftmost panel, cells were gated based on FSC-A and SSC-A profiles to find the population of cells. The middle panel gates cells by their FSC-A and FSC-H to isolate singlets. The final panel gates cells to isolate live cells from dead cells based on BV510 AquaZombie^™^ viability dye. BV510 negative cells were selected as the live cell population. b) Day 14 cell populations were compared to their FMO histograms for each marker of interest (left column), and the corresponding MFI was graphed (right column). All donors showed an increase in CD32B, CD14, and CD54 expression compared to the FMO group. c) Donor three cells thawed directly from the vial (top histogram) was compared to donor three cells after 14 days in culture (bottom histogram) for CD32B, CD14, and CD54. CD32B, CD14, and CD54 expression increased after being cultured for 14 days.

## Conclusion

In conclusion, (i) the LSECs used in this study are likely to have been isolated from zone 2 and 3 of the acinar lobule, (ii) co-culture of LSECs with hepatocytes on Liver-Chip maintains the characteristic LSEC morphology for 14 days, and (iii) Liver-Chip maintained the phenotypic expression of LSEC markers such as Stabilin 1, LYVE-1, CD32B, CD14 and CD54 for 14 Days, longer than shown for other published models. This study showed the potential of maintaining CD32B expression up to 14 days which is a relevant duration needed for therapeutic antibody discovery application. We also have shown the critical variables that directly affects the expression level of CD32B which is critical to mimic hepatic clearance of mAbs.

Therefore, this investigation shows both the versatility and the utility of the Liver-Chip allowing for future studies that could manipulate the expression of CD32B (e.g., to represent diseased liver) and other key markers expressed in liver cell lines. Overall, further optimization and validation of this model is required for use in the safety assessment of immuno-modulating FcγR binding biologics and in the prediction of pharmacokinetics/pharmacodynamics of biologics in humans. Thus, Liver-Chip offers a viable alternative to reduce or replace expensive and clinically poorly translatable *in vivo* models thereby paving the way for more predictive antibody discovery research with a strong impact on the 3Rs.

## Acknowledgements

We want to thank Daniel Rycroft for the helpful discussions around the need for new *in vitro* models in antibody drug discovery, Prof. Mark Cragg for providing AT10 antibody for CD32B expression analysis, Dr. Jason Ekert for contributions to developing new applications for Liver-Chip in large molecule discovery and Dr. Lorna Ewart for her critical review of the manuscript.

## Author contributions

PLC and SJ contributed equally to this paper. PLC and MH conceived the project. PLC, SJ and MH designed the study; PLC provided leadership for the whole project; SJ, JC, JV, SRJ contributed to chip seeding, maintenance, staining, sample preparation for SEM and chip sampling; AS and MK contributed to the flow cytometry data analysis; SJK and LE provided scientific oversight; PLC, SJ, JC, JV, MJ, AS contributed to writing the manuscript; PLC, MH, AS and LE critically reviewed the manuscript.

## Declarations

SJ, JC, JV, SRJ, MK, SJK and LE are employees of Emulate and may hold equity. PLC, MH, and JE are employees of GlaxoSmithKline and may hold equity. The human biological samples were sourced ethically, and their research use was in accord with the terms of the informed consents under an IRB/EC approved protocol.

## Data and materials availability

All analysed data and materials are available in the main text. Raw data supporting the findings of this study are available from the corresponding author upon reasonable request.

## Supplementary figure legend

**Supplementary Figure 1: Assessment of morphology and cell markers on the Liver-Chip.** a) Cells demonstrated typical morphology on day 3 in the top and bottom channels (top panel) at a flow rate of 30μl/h. By day 14, donors 2 and 4 showed a decline in cell number. b) Immunofluorescent staining images on day 14 – Donors 2 and 3 LSECs were positive for Stabilin-1, LYVE1, and CD32B, suggesting they were type 2 LSECs isolated from zones 2 and 3 of the acinar liver lobule. However, Donor 4 did not show CD32B or LYVE-1 expression, indicating there could be type 1 LSECs. c) Similar albumin levels were seen for all donors for the duration of the experiment, levelling out by day 14.

**Supplementary Figure 2: Criteria for Phenotypic morphology assessment on Chip**. Morphological assessment of cells based on the brightfield images are classified according to the matrix shown here a) for hepatocytes and b) for LSECs

**Supplementary Figure 3: Donor and passage information of LSECs**. We have used LSECs from 4 different donors with 4 different passage numbers to investigate the donor-to-donor variability and the passage number of the LSECs on the CD32B expression. *No additional donor information is available for this donor.

**Supplementary Figure 4: Representative immunofluorescence microscopy comparing CD32B, Stabilin-1, LYVE-1 and DAPI staining on 3 different LSEC donors on the Human Co-Culture Liver-Chip on Day 14.** From left to right, 3 Chips from each donor group were fixed and stained for LSECs markers such as CD32B, Stabilin-1 and LYVE-1. At Day 3 (a), Stabilin-1 expression is weak, LYVE-1 and CD32B are absent at Group 1, with moderate Stabilin-1 and low LYVE-1 expression on Group 2. Group 3 and 4 have strong expression of Stabilin-1 and LYVE-1 with moderate expression of CD32B.

On Day 14 (b), there is still no CD32b or LYVE-1 expression on Group 1, but homogenous Stabilin-1 expression was observed. LSECs in Groups 3 and 4 were positive for Stabilin-1, LYVE1, and CD32B (white arrows) on both timepoints suggesting they were type 2 LSECs isolated from zones 2 and 3 of the acinar liver lobule. However, fewer LSECs were found in Group 4 on Day 14.

**Supplementary Figure 5: Representative immunofluorescence microscopy comparing Stabilin-1 and CD32B antibodies on the Human Co-Culture Liver-Chip on Day 14.** Observed puncta believed to be staining artifacts (white arrows) seen in Ab1 CD32B antibody. Positive CD32B signal (pink arrows) seen in Ab2 (AT10 clone from University of Southampton Ab. Upcyte donor did not express CD32B until day 7 and maintained until day 14. Whereas the 0M and 1TB donors maintained CD32B expression from day 3 to day 14.

**Supplementary Figure 6: Experimental timeline for Chip culture of LSECs**. Hepatocytes are seeded to the top channel on day-2, and LSECs are seeded to the bottom channel on day 0. On days 3, 7, and 14, Chips were imaged for morphology, the effluent was collected for albumin assay, 3 Chips were fixed for IF imaging, and 1 Chip was fixed for SEM. On day 14, Flow cytometry analysis was done with n=3 Chips per group

## References

[1] M. W. Robinson, C. Harmon, and C. O’Farrelly, “Liver immunology and its role in inflammation and homeostasis,” Cellular & Molecular Immunology, vol. 13, no. 3, pp. 267–276, May 2016, doi: 10.1038/cmi.2016.3.

[2] I. N. Crispe, “The liver as a lymphoid organ.,” Annual review of immunology, vol. 27, pp. 147–63, 2009, doi: 10.1146/annurev.immunol.021908.132629.

[3] L. D. DeLeve and A. C. Maretti-Mira, “Liver Sinusoidal Endothelial Cell: An Update.,” Seminars in liver disease, vol. 37, no. 4, pp. 377–387, 2017, doi: 10.1055/s-0037-1617455.

[4] K. B. Halpern et al., “Paired-cell sequencing enables spatial gene expression mapping of liver endothelial cells.,” Nature biotechnology, vol. 36, no. 10, pp. 962–970, 2018, doi: 10.1038/nbt.4231.

[5] S. A. MacParland et al., “Single cell RNA sequencing of human liver reveals distinct intrahepatic macrophage populations.,” Nature communications, vol. 9, no. 1, p. 4383, 2018, doi: 10.1038/s41467-018-06318-7.

[6] J. Poisson et al., “Liver sinusoidal endothelial cells: Physiology and role in liver diseases.,” Journal of hepatology, vol. 66, no. 1, pp. 212–227, 2017, doi: 10.1016/j.jhep.2016.07.009.

[7] E. Wisse, R. B. de Zanger, K. Charels, P. van der Smissen, and R. S. McCuskey, “The liver sieve: considerations concerning the structure and function of endothelial fenestrae, the sinusoidal wall and the space of Disse.,” Hepatology (Baltimore, Md.), vol. 5, no. 4, pp. 683–92, doi: 10.1002/hep.1840050427.

[8] K. K. Sørensen, J. Simon-Santamaria, R. S. McCuskey, and B. Smedsrød, “Liver Sinusoidal Endothelial Cells.,” Comprehensive Physiology, vol. 5, no. 4, pp. 1751–74, Sep. 2015, doi: 10.1002/cphy.c140078.

[9] L. D. DeLeve, “Liver sinusoidal endothelial cells and liver regeneration.,” The Journal of clinical investigation, vol. 123, no. 5, pp. 1861–6, May 2013, doi: 10.1172/JCI66025.

[10] J. C. Anania, A. M. Chenoweth, B. D. Wines, and P. M. Hogarth, “The Human FcγRII (CD32) Family of Leukocyte FcR in Health and Disease.,” Frontiers in immunology, vol. 10, p. 464, 2019, doi: 10.3389/fimmu.2019.00464.

[11] Y. Xu et al., “Fc gamma Rs modulate cytotoxicity of anti-Fas antibodies: implications for agonistic antibody-based therapeutics.,” Journal of immunology (Baltimore, Md.□: 1950), vol. 171, no. 2, pp. 562–8, Jul. 2003, doi: 10.4049/jimmunol.171.2.562.

[12] F. Li and J. v. Ravetch, “Apoptotic and antitumor activity of death receptor antibodies require inhibitory Fc receptor engagement,” Proceedings of the National Academy of Sciences, vol. 109, no. 27, pp. 10966–10971, Jul. 2012, doi: 10.1073/pnas.1208698109.

[13] L. P. Ganesan et al., “FcγRIIb on Liver Sinusoidal Endothelium Clears Small Immune Complexes,” The Journal of Immunology, vol. 189, no. 10, pp. 4981–4988, Nov. 2012, doi: 10.4049/jimmunol.1202017.

[14] S. Khan et al., “The Association for Human Pharmacology in the Pharmaceutical Industry London Meeting October 2019: Impending Change, Innovation, and Future Challenges,” Frontiers in Pharmacology, vol. 11, Nov. 2020, doi: 10.3389/fphar.2020.580560.

[15] A. L. White et al., “Interaction with FcγRIIB Is Critical for the Agonistic Activity of Anti-CD40 Monoclonal Antibody” The Journal of Immunology, vol. 187, no. 4, pp. 1754–1763, Aug. 2011, doi: 10.4049/jimmunol.1101135.

[16] F. Li and J. v Ravetch, “Inhibitory Fcγ receptor engagement drives adjuvant and antitumor activities of agonistic CD40 antibodies.,” Science (New York, N.Y.), vol. 333, no. 6045, pp. 1030–4, Aug. 2011, doi: 10.1126/science.1206954.

[17] F. R. Brennan et al., “Safety testing of monoclonal antibodies in non-human primates: Case studies highlighting their impact on human risk assessment,” mAbs, vol. 10, no. 1, pp. 1–17, Jan. 2018, doi: 10.1080/19420862.2017.1389364.

[18] N. H. Segal et al., “Results from an Integrated Safety Analysis of Urelumab, an Agonist Anti-CD137 Monoclonal Antibody.,” Clinical cancer research□: an official journal of the American Association for Cancer Research, vol. 23, no. 8, pp. 1929–1936, 2017, doi: 10.1158/1078-0432.CCR-16-1272.

[19] S. B. Laffan et al., “Immune complex disease in a chronic monkey study with a humanised, therapeutic antibody against CCL20 is associated with complement-containing drug aggregates,” PLOS ONE, vol. 15, no. 4, p. e0231655, Apr. 2020, doi: 10.1371/journal.pone.0231655.

[20] N. Stuurman and J. R. Swedlow, “Software Tools, Data Structures, and Interfaces for Microscope Imaging,” Cold Spring Harbor Protocols, vol. 2012, no. 1, p. pdb.top067504, Jan. 2012, doi: 10.1101/pdb.top067504.

[21] P. Shah, V. Sundaram, and E. Björnsson, “Biologic and Checkpoint Inhibitor-Induced Liver Injury: A Systematic Literature Review.,” Hepatology communications, vol. 4, no. 2, pp. 172–184, Feb. 2020, doi: 10.1002/hep4.1465.

[22] Y. Koui et al., “An In Vitro Human Liver Model by iPSC-Derived Parenchymal and Non-parenchymal Cells,” Stem Cell Reports, vol. 9, no. 2, pp. 490–498, Aug. 2017, doi: 10.1016/j.stemcr.2017.06.010.

[23] S. D. Ramachandran et al., “In Vitro Generation of Functional Liver Organoid-Like Structures Using Adult Human Cells.,” PloS one, vol. 10, no. 10, p. e0139345, 2015, doi: 10.1371/journal.pone.0139345.

[24] Lorna Ewart et al., “Qualifying a human Liver-Chip for predictive toxicology: Performance assessment and economic implications”, Accessed: Jan. 13, 2022. [Online]. Available: https://www.biorxiv.org/content/10.1101/2021.12.14.472674v3

[25] K.-J. Jang et al., “Reproducing human and cross-species drug toxicities using a Liver-Chip,” Science Translational Medicine, vol. 11, no. 517, Nov. 2019, doi: 10.1126/scitranslmed.aax5516.

[26] L. Prodanov et al., “Long-term maintenance of a microfluidic 3D human liver sinusoid,” Biotechnology and Bioengineering, vol. 113, no. 1, pp. 241–246, Jan. 2016, doi: 10.1002/bit.25700.

[27] T. Kostrzewski et al., “A Microphysiological System for Studying Nonalcoholic Steatohepatitis,” Hepatology Communications, vol. 4, no. 1, pp. 77–91, Jan. 2020, doi: 10.1002/hep4.1450.

[28] S. S. Bale eř al., “In vitro platforms for evaluating liver toxicity,” Experimental Biology and Medicine, vol. 239, no. 9, pp. 1180–1191, Sep. 2014, doi: 10.1177/1535370214531872.

[29] A. R. Baudy et al., “Liver microphysiological systems development guidelines for safety risk assessment in the pharmaceutical industry” Lab on a Chip, vol. 20, no. 2, pp. 215–225, 2020, doi: 10.1039/C9LC00768G.

[30] B. R. Ware, M. J. Durham, C. P. Monckton, and S. R. Khetani, “A Cell Culture Platform to Maintain Long-term Phenotype of Primary Human Hepatocytes and Endothelial Cells.,” Cellular and molecular gastroenterology and hepatology, vol. 5, no. 3, pp. 187–207, Mar. 2018, doi: 10.1016/j.jcmgh.2017.11.007.

[31] S. March, E. E. Hui, G. H. Underhill, S. Khetani, and S. N. Bhatia, “Microenvironmental regulation of the sinusoidal endothelial cell phenotype in vitro,” Hepatology, vol. 50, no. 3, pp. 920–928, Sep. 2009, doi: 10.1002/hep.23085.

[32] K. Szafranska, L. D. Kruse, C. F. Holte, P. McCourt, and B. Zapotoczny, “The wHole Story About Fenestrations in LSEC,” Frontiers in Physiology, vol. 12, Sep. 2021, doi: 10.3389/fphys.2021.735573.

[33] E. Pandey, A. S. Nour, and E. N. Harris, “Prominent Receptors of Liver Sinusoidal Endothelial Cells in Liver Homeostasis and Disease,” Frontiers in Physiology, vol. 11, Jul. 2020, doi: 10.3389/fphys.2020.00873.

[34] K. Szafranska, L. D. Kruse, C. F. Holte, P. McCourt, and B. Zapotoczny, “The wHole Story About Fenestrations in LSEC.,” Frontiers in physiology, vol. 12, p. 735573, 2021, doi: 10.3389/fphys.2021.735573.

[35] V. C. Cogger, N. J. Hunt, and D. G. le Couteur, “Fenestrations in the Liver Sinusoidal Endothelial Cell,” in The Liver, Wiley, 2020, pp. 435–443. doi: 10.1002/9781119436812.ch35.

[36] Y. Du et al., “Mimicking liver sinusoidal structures and functions using a 3D-configured microfluidic chip,” Lab on a Chip, vol. 17, no. 5, pp. 782–794, 2017, doi: 10.1039/C6LC01374K.

[37] S. R. Khetani and S. N. Bhatia, “Microscale culture of human liver cells for drug development,” Nature Biotechnology, vol. 26, no. 1, pp. 120–126, Jan. 2008, doi: 10.1038/nbt1361.

[38] S.-Y. Chang et al., “Characterization of rat or human hepatocytes cultured in microphysiological systems (MPS) to identify hepatotoxicity.,” Toxicology in vitro□: an international journal published in association with BIBRA, vol. 40, pp. 170–183, Apr. 2017, doi: 10.1016/j.tiv.2017.01.007.

[39] J. LeCouter et al., “Angiogenesis-Independent Endothelial Protection of Liver: Role of VEGFR-1,” Science, vol. 299, no. 5608, pp. 890–893, Feb. 2003, doi: 10.1126/science.1079562.

[40] Niikolett Nagy et al., “in vitro organoid model of the human liver sinusoid.” [Online]. Available: https://www.upcyte.com/wp-content/uploads/2020/04/28Poster_uptec_201906_LSEC_chip.pdf

[41] M. E. Fomin, A. I. Beyer, and M. O. Muench, “Human fetal liver cultures support multiple cell lineages that can engraft immunodeficient mice,” Open Biology, vol. 7, no. 12, p. 170108, Dec. 2017, doi: 10.1098/rsob.170108.

[42] A. Karrar, U. Broomé, M. Uzunel, A. R. Qureshi, and S. Sumitran-Holgersson, “Human liver sinusoidal endothelial cells induce apoptosis in activated T cells: a role in tolerance induction.,” Gut, vol. 56, no. 2, pp. 243–52, Feb. 2007, doi: 10.1136/gut.2006.093906.

