## Supplemental Figures for "Phenotypic Characterization of Liver Sinusoidal Endothelial Cells on the Human Liver-Chip for Potential *in vitro* Therapeutic Antibody Pharmacology Applications"

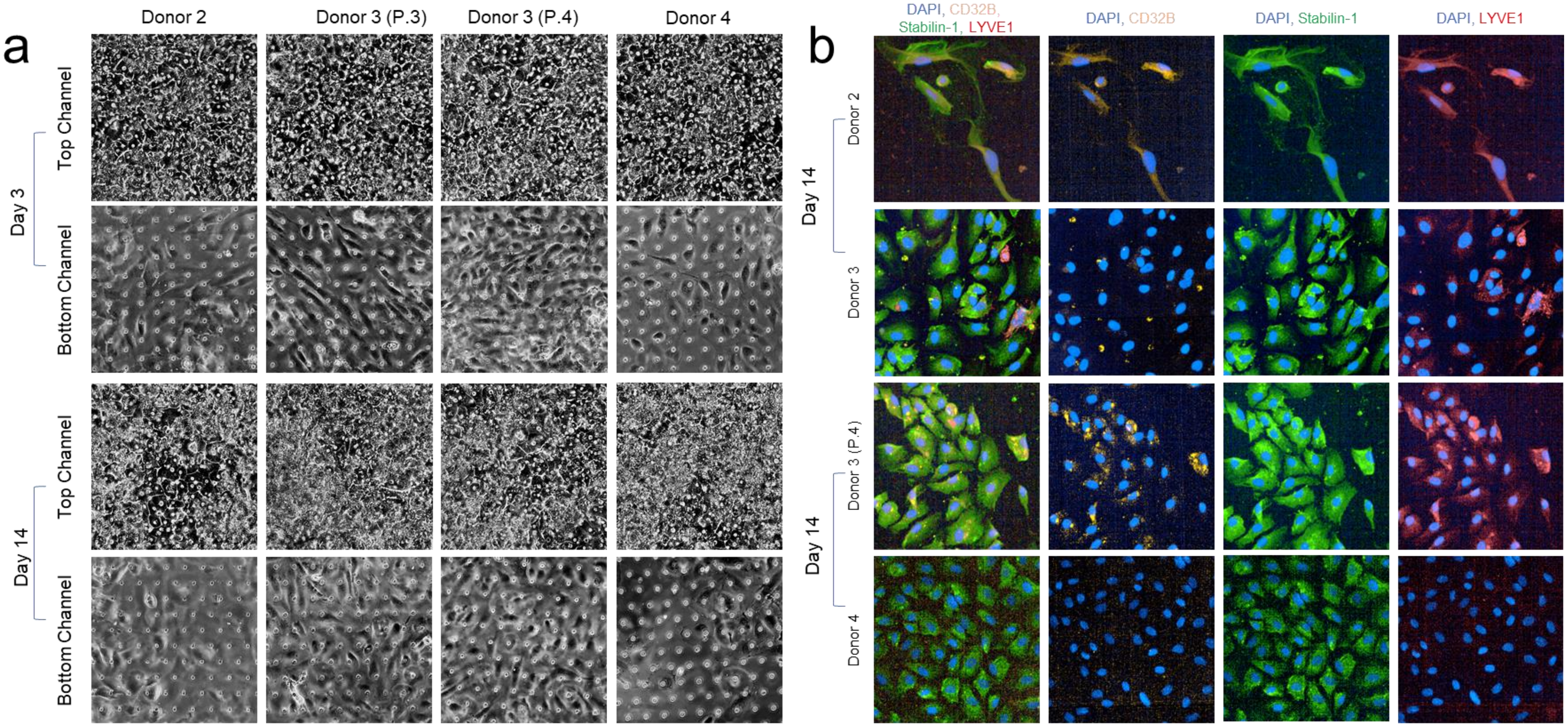


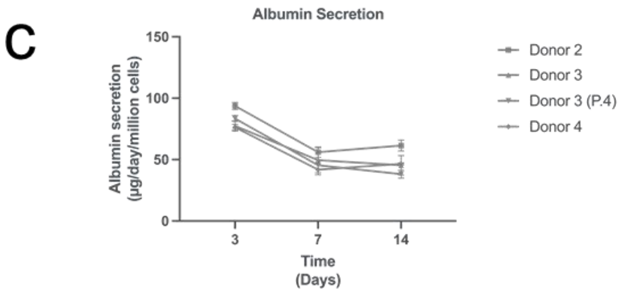


**Supplementary Figure 1: Assessment of morphology, and cell markers on the Liver-Chip.** a) Cells demonstrated typical morphology on day 3 in both the top and bottom channels (top panel) at a flow rate of 30µl/h. By day 14 donors 2 and 4 showed a decline in cell number. b) Immunofluorescent staining images on day 14 – Donors 2 and 3 LSECs were positive for Stabilin-1, LYVE1, and CD32B, suggesting they were type 2 LSECs isolated from zones 2 and 3 of the acinar liver lobule. However, Donor 4 did not show CD32b or LYVE-1 expression on, indicating these could be type 1 LSECs. c) Similar albumin levels were seen for all donors for the duration of the experiment, leveling out by day 14.

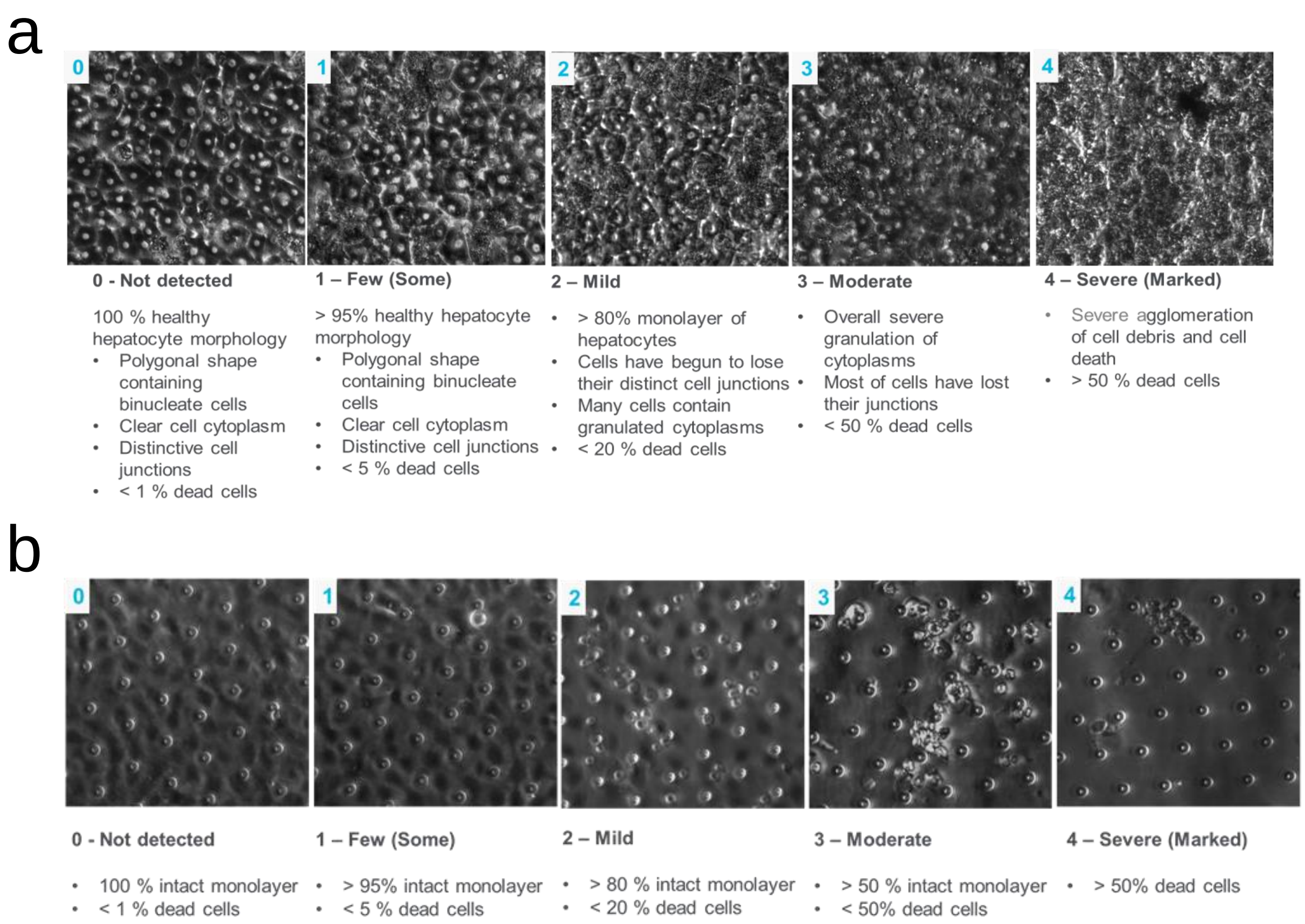


**Supplementary Figure 2: Criteria for Phenotypic morphology assessment on Chip**. Morphological assessment of cells based on the brightfield images are classified according to the matrix shown here a) for hepatocytes and **b**) for LSECs

| Donor Number | Vendor | Lot Information | Passage Number | Donor information |
|---|---|---|---|---|
| Donor 1 | Upcyte | 462-20210325.1 | Passage unknown | Female, 62, Caucasian |
| Donor 2 | Cell Systems | 566.01.01.01.1TB | Passage 3 | Female, 27, Caucasian |
| Donor 3 | Cell Systems | 566.02.02.05.0M | Passage 3 | Male* |
| Donor 4 | iXCells | 200356 | Passage 2 | Male, 55, Caucasian |

**Supplementary Figure 3: Donor and passage information of LSECs**. LSECs from 4 donors with 3 different passage numbers were used to investigate the effect of donor-to-donor variability on the CD32b expression. *Additional donor information is not available for this donor.

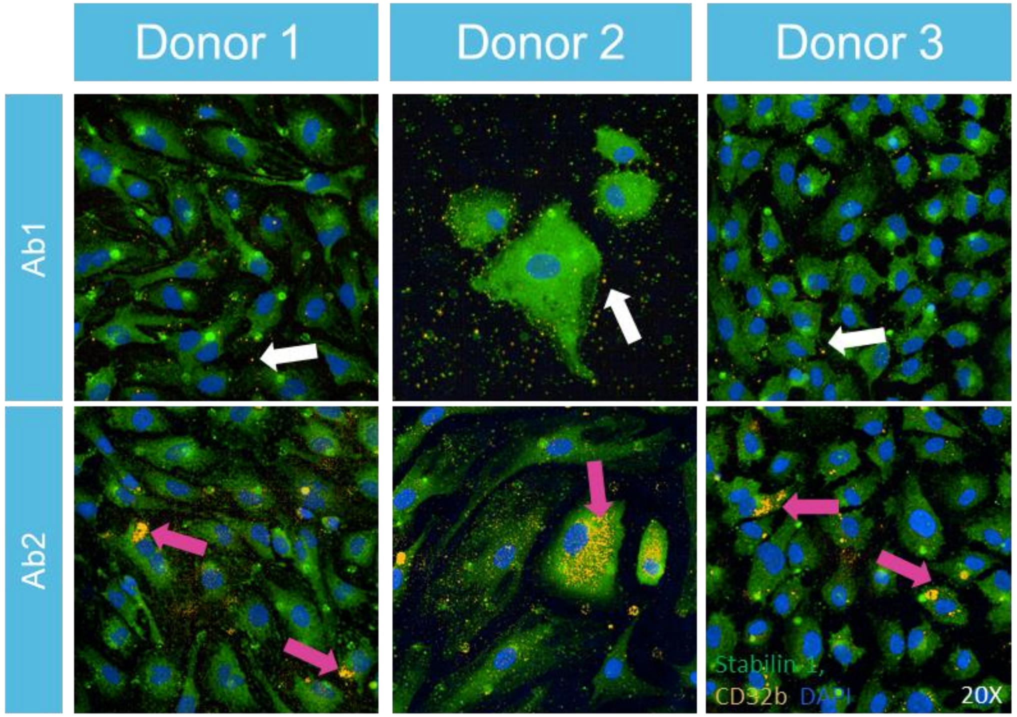


**Supplementary Figure 4: Representative immunofluorescence images comparing Stabilin-1 and CD32b antibodies on the Human Co-Culture Liver-Chip on Day 14.** LSECs stained with Ab1 (Mouse myeloma cell line NS0-derived recombinant human Fc gamma RII/CD32 Ala46-Pro217) CD32b Ab were Stabilin $1^{+}$ $CD32B^{-}$ while, LSECs stained with CD32b Ab2 (AT10 clone from University of Southampton Ab) were Stabilin$1^{+}$ $CD32B^{+}$.

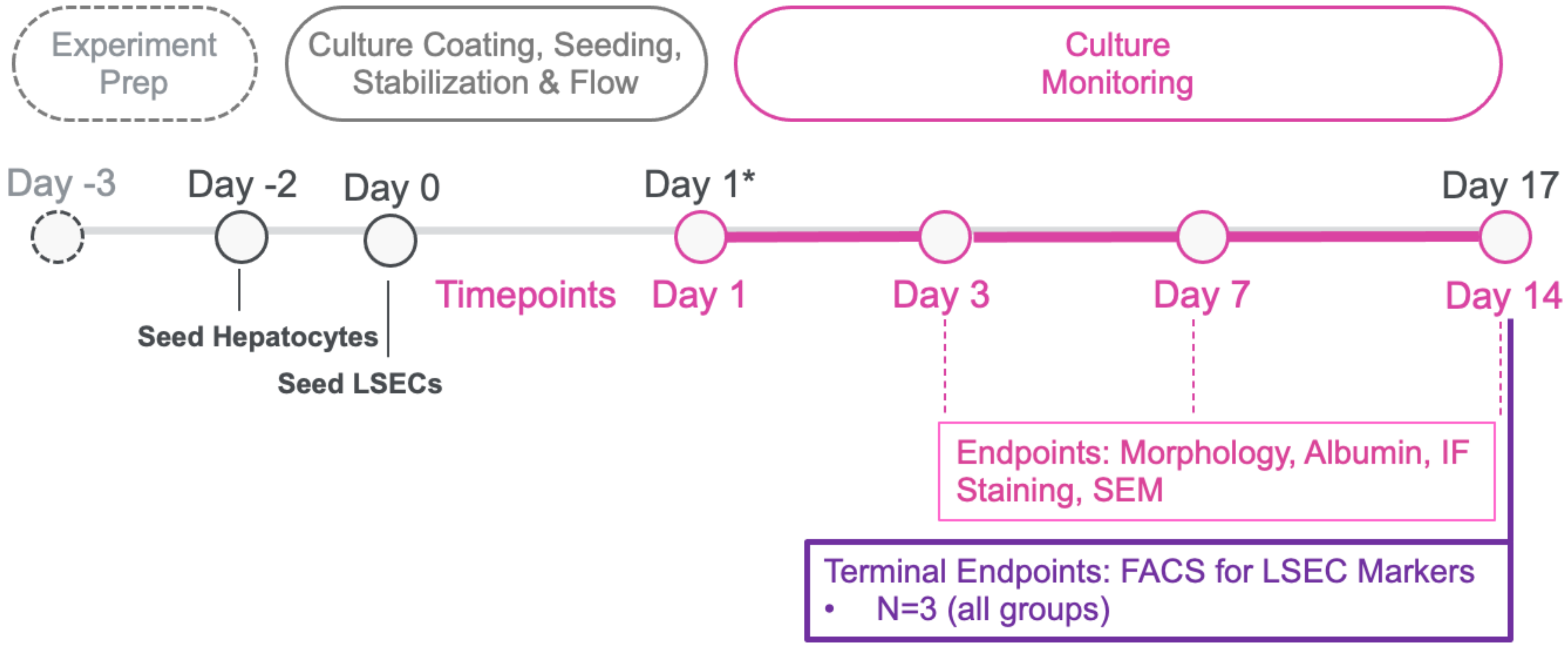


**Supplementary Figure 5: Experimental timeline for Chip culture of LSECs**. Hepatocytes are seeded to the Top Channel on day -2 and LSECs are seeded to the Bottom Channel on day 0. On days 3, 7, and 14, Chips were imaged for morphology, the effluent was collected for albumin assay, 3 Chips were fixed for IF imaging, and 1 Chip was fixed for SEM. On day 14 Flow cytometry analysis was done with n=3 Chips per group

| Antibody | Vendor | Clone |
|---|---|---|
| **Zombie Aqua** | Biolegend | - |
| **CD32B** | University of Southampton | AT10 |
| **CD54** | Biolegend | HA58 |
| **CD14** | Biolegend | 63D3 |

**Supplementary Figure 6: Vendor and Clone information for Flow Cytometry**. Cells from donors 1-3 were stained with a Live/Dead stain (Zombie Aqua) and three additional antibodies to confirm marker expression.
